# Protein-free division of giant unilamellar vesicles controlled by enzymatic activity

**DOI:** 10.1101/2019.12.30.881557

**Authors:** Yannik Dreher, Joachim P. Spatz, Kerstin Göpfrich

## Abstract

Cell division is one of the hallmarks of life. Success in the bottom-up assembly of synthetic cells will, no doubt, depend on strategies for the controlled autonomous division of protocellular compartments. Here, we describe the protein-free division of giant unilamellar lipid vesicles (GUVs) based on the combination of two physical principles – phase separation and osmosis. We visualize the division process with confocal fluorescence microscopy and derive a conceptual model based on the vesicle geometry. The model successfully predicts the shape transformations over time as well as the time point of the final pinching of the daughter vesicles. Remarkably, we show that two fundamentally distinct yet highly abundant processes – water evaporation and metabolic activity – can both regulate the autonomous division of GUVs. Our work may hint towards mechanisms that governed the division of protocells and adds to the strategic toolbox of bottom-up synthetic biology with its vision of bringing matter to life.

---

*”Omni cellulae es cellulae.”* From the point of view of modern science, Raspail’s realization from 1825 [1], popularized by Virchow [2], may state the obvious: Every living cell found on Earth today originates from a preexisting living cell. Bottom-up synthetic biology, however, is challenging this paradigm with the vision to create a synthetic cell from scratch [3, 4]. Success unquestionably entails that the synthetic cells must have the capacity to produce offspring, making the implementation of synthetic cell division an exciting goal [5, 6, 7, 8]. Over the course of evolution, living cells have developed a sophisticated machinery to divide their compartments in a highly regulated manner. The reconstitution of a minimal set of components from the procaryotic divisome seems to be a promising route towards synthetic cell division. FtsZ [9, 6], ESCRT [10, 11] and Min proteins [12] led to shape transformations in lipid vesicles, including budding [11] and the formation of tight constrictions [6]. However, reproducible minimal cell division based on common biological mechanisms has not yet been achieved. These challenges leave room for creative approaches, seeking solutions beyond the mimicry of today’s biological cells. One exciting strategy is to assemble a division machinery de novo, by designing active, not necessarily protein-based nanomachines. DNA origami structures have been used to shape and remodel lipid vesicles [13, 14, 15], although active force-generating contractile motors remain a distant goal. A shortcut towards synthetic cell division is the mechanical division of liposomes, which was achieved with microfluidic splitters [16]. While this is not autonomous, it may jump start exciting new directions by circumventing a longstanding challenge. The exploitation of physico-chemical mechanisms, on the other hand, could lead to autonomous division. Noteworthy theoretical and experimental work describes the shape transformations of single-component [17, 18, 19] as well as phase-separated liposomes [20, 21], determined by the minimum elastic energy for a given surface-to-volume ratio. Within this phase space, two vesicles connected with a tight neck resemble division most closely [18]. However, neck fission does normally not occur. There are only few examples of complete budding or splitting in the literature. The earliest one describes small buds dissociating from a parent vesicle [22]; later work realizing division into more equally sized compartments reports occasional observations [23], relies on multilamellar vesicles [24] or liquid-liquid phase separation [25].

Here, we demonstrate full spatiotemporal control over the autonomous and protein-free division of GUVs. The division is based on a combination of two physical mechanisms, namely phase separation and osmosis. This allows us to precisely predict the division process and its outcome based on geometrical considerations. To the best of our knowledge, we demonstrate for the first time that the division of GUVs can be regulated by metabolic activity. Our results prompt to ask whether similar mechanisms may have sustained cell division at the onset of live [26, 27]. Similarly, remnants thereof may still play a role in cell biology today, e.g. in the generation of intracellular vesicles or possibly to support membrane curvature during the division process [28, 29, 30].

## Results

### GUV division induced by phase separation and osmotic pressure

GUVs – i.e. micron-sized vesicles enclosed by a single lipid bilayer – are the most commonly used compartment type for the assembly of synthetic cells [4]. To realize a simple physical model for their division, we assume that two steps have to be implemented experimentally: Step 1) Define the plane of division; Step 2) Increase the surface-to-volume ratio to allow for the formation of two smaller daughter compartments from a single large compartment. To realize Step 1, we choose lipid phase separation as a strategy to obtain the required symmetry break. This allows us to define the plane of division as the interface of the liquid disordered (ld, orange) and the liquid ordered (lo, green) phase as illustrated in Figure 1**a**. Note that the number of lipids in the membrane, i.e. the surface area of the GUV, remains constant. Hence an increase in surface-to-volume ratio (Step 2) requires a reduction of the GUV’s inner volume. To this end, we rely on osmosis. In the simplest case, water evaporation leads to an increase of the osmolarity, i.e. the concentration of solutes in aqueous solution, outside the GUVs. This, in turn, causes near instantaneous water efflux through the GUV membrane [31] as illustrated in Figure 1**b**. Minimizing the line tension at the phase boundary and increasing the curvature, the GUV deforms to store the excess membrane area caused by the volume reduction. This can ultimately lead to division: Although the final pinching of the lipid constriction comes with an energy cost for opening up the bilayer structure, it also leads to complete demixing of the lipids, removing the line tension at the phase boundary [32, 21]. Therefore – unlike in the case of single-phase GUVs – complete division could result in a net energy gain.

**Figure 1:**
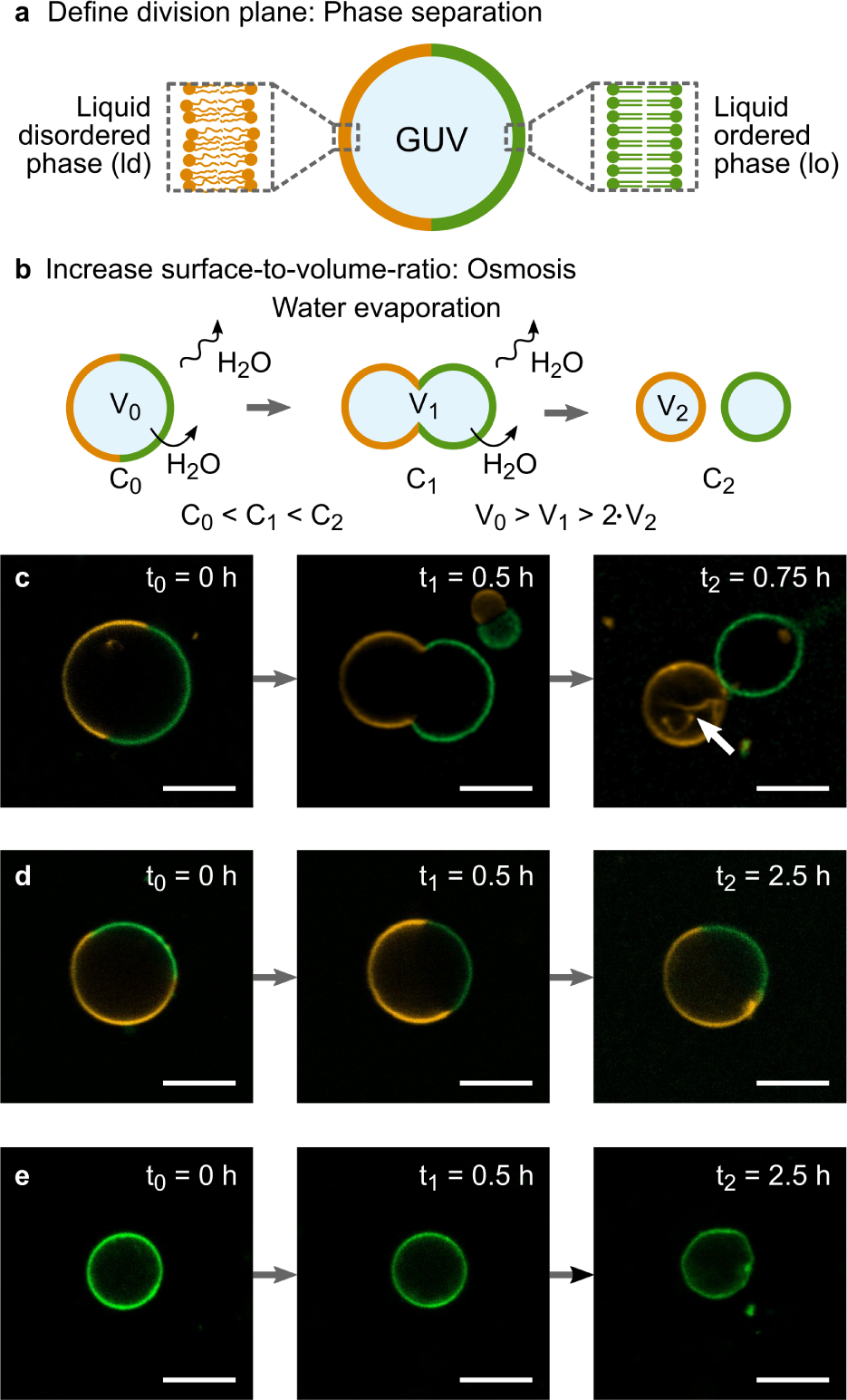
Osmotic pressure-driven division of phase-separated GUVs. Schematic illustration of the division mechanism relying on **a** phase separation of the GUVs and **b** osmosis induced by water evaporation. C_*i*_ denotes the osmolarity outside of the GUVs and V_*i*_ describes their volume; *i* = 0, 1, 2. **c** Time series of confocal fluorescence images of the division process of a phase-separated GUV upon water evaporation. The white arrow highlights the formation of lipid tubes after the division is completed. **d** Control experiment with a phase-separated GUV in a sealed observation chamber preventing water evaporation. **e** Control experiment with a single-phase GUV in an unsealed observation chamber allowing for water evaporation. LissRhod PE labeled the ld phase (orange, *λ*_ex_ = 561 nm) and 6-FAM-labeled cholesterol-tagged DNA partitioned in the lo phase (green, *λ*_ex_ = 488 nm). Single-phase GUVs were labeled via Atto488-DOPE (green, *λ*_ex_ = 488 nm). Scale bars: 10 µm.

To implement the proposed division mechanism experimentally, we initially need to accomplish the formation of a GUV composed of two distinct hemispheres. At room temperature, a lipid mixture composed of 27.125 % DOPC, 24.75 % cholesterol, 37.125 % DPPC, 10 % CL and 1 % LissRhod PE (for visualization) separates into a liquid ordered (lo, saturated lipids) phase and a liquid disordered (ld, unsaturated lipids) phase [33]. With confocal fluorescence microsocopy, we confirm the successful formation of phase-separated GUVs as shown in Figure 1**c** (left). LissRhod PE partitions into the ld phase (orange), whereas cholesterol-tagged 6-FAM-labeled DNA self-assembles selectively into the lo phase (green). Note that the self-assembly of the cholesterol-tagged DNA into the lipid bilayer requires Mg^2+^ in the buffer (see Supplementary Figure S1). We increase the osmolarity outside the GUVs by allowing for continuous water evaporation during the imaging process simply by observing the GUVs in an unsealed observation chamber at room temperature. Figure 1**c** shows a time series of confocal fluorescence images taken over the course of 75 minutes. Remarkably, we observe the formation of a constriction at the interface of the two phases, eventually leading to complete division. Note that GUVs often remain in close contact although they are fully divided due to electrostatic interactions mediated by the Mg^2+^ ions in the buffer. Only after soft shaking, they are found in complete isolation (see Supplementary Figure S2). If Mg^2+^ is removed from the buffer, the GUVs separate as soon as the division is completed (see also Figure 4**d**). This is the ultimate proof that complete neck scission occurred. It also shows that the membrane-bound DNA is not a key factor aiding the division, it was merely used as a fluorescent label. After the division process is completed, lipid tubulation can occur (highlighted by the white arrow in Figure 1**c**, right) to store excess membrane area resulting from further deflation of the compartments.

To confirm that the demonstrated division mechanism requires both, lipid phase separation and osmosis, we performed two sets of control experiments: First, the phase-separated vesicles are placed in a sealed observation chamber which does not allow for evaporation of water. As expected for constant osmolarity, no deformation is visible even after 2.5 h (see Figure 1**d**). Second, in order to show that increasing the surface-to-volume ratio of GUVs alone is not sufficient to achieve division, we expose single-phase GUVs to an osmolarity gradient by water evaporation (see Figure 1**e**). The evaporation of water leads to deflation of the GUV, resulting in increased membrane fluctuations or lipid tube formation to store the excess membrane area. However, this effect alone does not lead to division. From these experiments, we can deduce a well-defined physical mechanism for the replication of protocells based on phase separation and osmosis. Since evaporation is a common phenomenon in nature, it is conceivable that early protocells were able to exploit this effect when phospholipids emerged and led to phase separation in fatty acid membranes [26, 27]. Similar mechanisms may still play a role: The fact that metabolic activity drives phase separation in the endoplasmatic reticulum [34] may hint that phase separation could be relevant for the generation of intracellular vesicles. Genetically modified bacteria have been shown to divide without the relevant division machinery [35]. Moreover, it seems possible that lymphocyte proliferation rates increase under hypertonic conditions [36].

### Theoretical prediction of the division process

Next, we set out to predict the osmolarity change required to achieve division of a phase-separated GUV. For this purpose, we develop a geometrical model describing the shape of the GUV throughout the deformation process as two spherical caps with a base radius *s*_0_ for the initially spherical GUV and *s < s*_0_ for the deformed GUV. One of them represents the ld phase with a surface area *A*_ld_ and the other one the lo phase with a surface area *A*_lo_, respectively. This representation provides a good approximation of our experimentally observed as well as the theoretically derived [20, 21] GUV shapes. Figure 2**a** indicates the relevant geometrical properties of the GUV, which can be extracted from confocal images. While living cells normally undergo symmetric division, where both daughter compartments are of similar size, some processes like oocyte maturation rely on assymmetric division [37]. In our model system, *A*_lo_ = *A*_ld_ leads to symmetric division as already demonstrated experimentally, whereas asymmetric division should be achieved otherwise. We assume that the total area *A*_tot_ remains constant throughout the division process as the number of lipids in the membrane does not change. If, however, the outer osmolarity increases due to water evaporation or addition of solutes (*C > C*_0_), the volume of the GUV will decrease due to water efflux. This nearly instantaneous process rapidly equilibrates the outer and the inner osmolarity [31]. The reduced inner volume is then given by 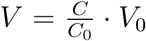. The resulting excess membrane area allows for deformation of the initially spherical GUV. As theoretically predicted [20], deformation minimizes the phase boundary (*s < s*_0_) to reduce the line tension between the two lipid phases. To quantify the progression of the division process, we define a division parameter *d*:

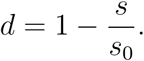

*d* is 0 for the initial spherical GUV and 1 for a divided GUV. Based on these geometrical considerations, the osmolarity ratio *C/C*_0_ needed to achieve a certain amount of deformation *d* for a symmetric GUV (*A*_ld_ = *A*_lo_) can be calculated as

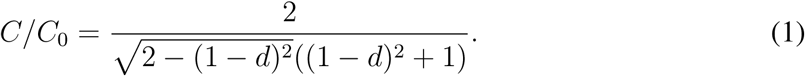

**Figure 2:**
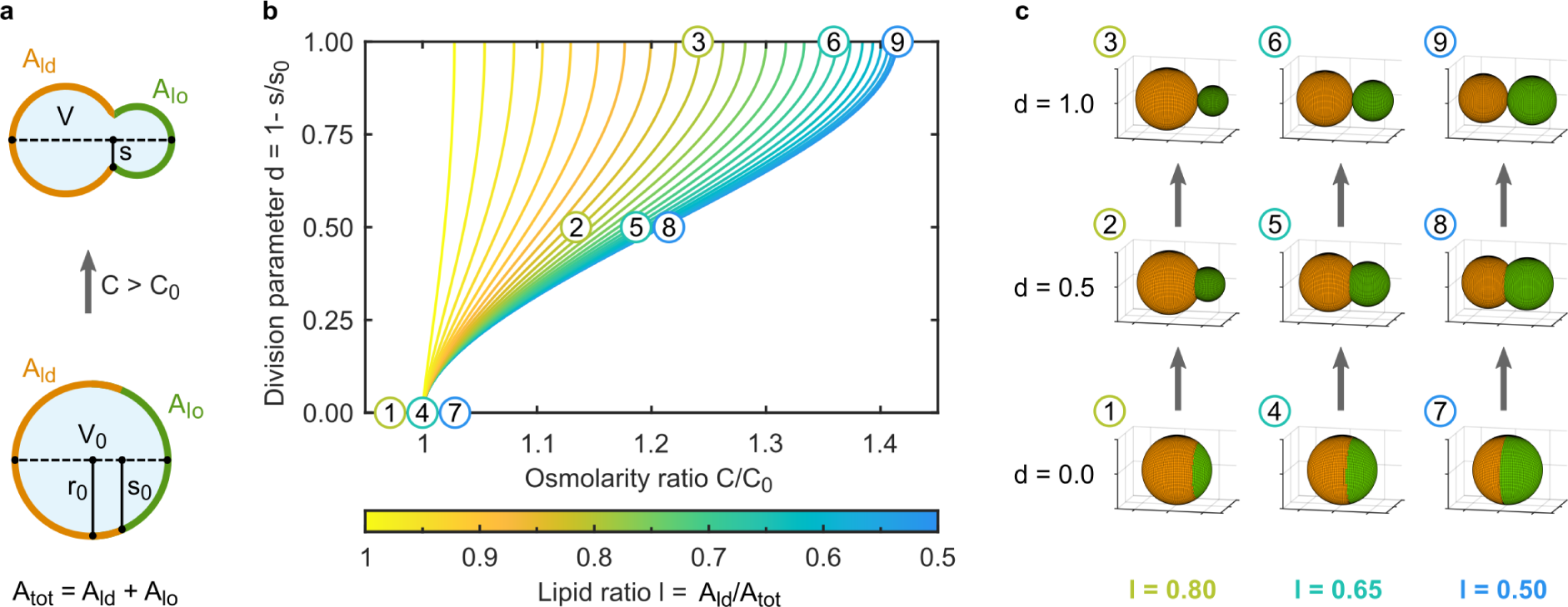
Theoretical predictions for the division process of phase-separated GUVs based on an increase in the osmolarity ratio. **a** Schematic describing the relevant geometrical properties of a deformed GUV (top) and its initially spherical state (bottom). *A*_ld_ and *A*_lo_ are the surface areas of the spherical caps representing the two phases. *s*_0_ is the radius of the base of the caps, *V*_0_ the volume and *r*_0_ the radius of the initially spherical GUV. *s* is the reduced radius of the base of the caps and *V* the reduced volume of the deformed GUV. **b** Theoretical prediction of the division parameter *d* as a function of osmolarity ratio *C/C*_0_ for different lipid ratios *l*. *d* = 0 corresponds to a spherical GUV, *d* = 1 to a fully divided one. **c** Predicted shapes of GUVs with different lipid ratios (*l* = 0.80, 0.65, 0.5) at defined points during the division process (*d* = 0.0, 0.5, 1.0). The corresponding positions are indicated in the plot in **b**.

For the more general case of asymmetric GUVs with *A*_ld_ ≠ *A*_lo_ we define a lipid ratio parameter *l* with 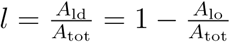. We hence obtain

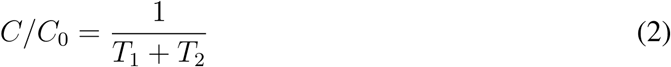

with 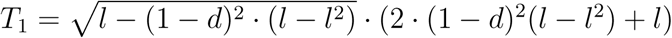 and

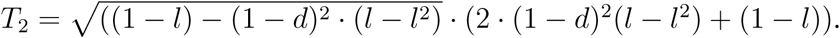

This geometrical model makes three predictions which we can test experimentally: First of all, it is important to note that both Equation 1 and 2 depend only on the division parameter *d* and the lipid ratio parameter *l* but not on the initial radius *r*_0_ of the GUV. Thus, we postulate that the osmolarity ratio required for division is independent of the size of the GUV. Second, we can calculate the osmolarity ratio required to divide a GUV: For *d* = 1 (corresponding to a fully divided GUV) we obtain 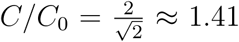 for a symmetric GUV according to Equation 1. Third, according to Equation 2, GUVs with higher asymmetry should require lower osmolarity ratios for complete division. Figure 2**b** shows the predicted division parameter *d* as a function of the osmolarity ratio *C/C*_0_ for different lipid ratios *l*. Note that a GUV with *l* = 0.8 divides already at an osmolarity ratio of approximately 1.22 (compared 1.41 for symmetric GUVs with *l* = 0.5). For clarity, Figure 2**c** displays the predicted shapes of the GUVs corresponding to specific points as indicated in Figure 2**b**. Most interestingly, any process that changes the osmolarity ratio should, in theory, lead to division – suggesting that metabolic activity could drive and regulate the division process.

### Quantitative comparison of experiments and theoretical predictions

To test the predictions of our model in a quantitative manner, we observe symmetric GUVs (*l* = 0.5) at different well-defined osmolarity ratios *C/C*_0_. For this purpose, phase-separated GUVs were immersed in buffer solutions with an increased osmolarity and sealed in an observation chamber to prevent evaporation. It is crucial to immerse the GUVs slowly to avoid tubulation of the GUVs (see Supplementary Figures S3, S4). Figure 3**a** shows the theoretically predicted shapes for the different osmolarity ratios; corresponding representative confocal fluorescence images are presented in Figure 3**b**. Note that the shapes are static since the osmolarity ratio is kept constant, unlike in the case of evaporation. This allows us to extract the geometrical parameters required to calculate the division parameter *d* from multiple images. Histograms of the distribution of *d* at a given osmolarity ratio are shown in Figure 3**c**. To verify the predicted size independence of the division process, we used the images of the deformed GUVs to calculate the radius *r*_0_ of the initially spherical GUV. The scatter plot of *d* over *r*_0_ is shown in Figure 3**d**. As expected, no significant size-dependent deviations from the theoretical value (blue line) can be observed. Note that effects like the vesicle size-dependent bending energy have been neglected in our model, which is validated by our experimental results. For small unilamellar vesicles below 1 *µ*m in diameter, the bending energy will dominate the line tension and size effects come into play [22]. As a quantitative comparison of the experimental results with the theoretical predictions (Equation 1), we plot the mean division parameter as a function of the osmolarity ratio. Figure 3**e** shows that the experimental data agrees well with the theoretical prediction (solid blue line). Deviations may occur due to the fact that GUVs are imaged in solution and can hence rotate in the confocal plane. When trapping them on a surface or in a microfluidic trapping device, interfacial effects can lead to lipid tubulation and hinder the division process (see Supplementary Figure S5). Furthermore, the predicted shape itself is just an approximation of the actual GUV. It is remarkable that a simple conceptual model based on only few geometrical parameters could quantitatively predict the division behaviour of cell-sized lipid vesicles.

**Figure 3:**
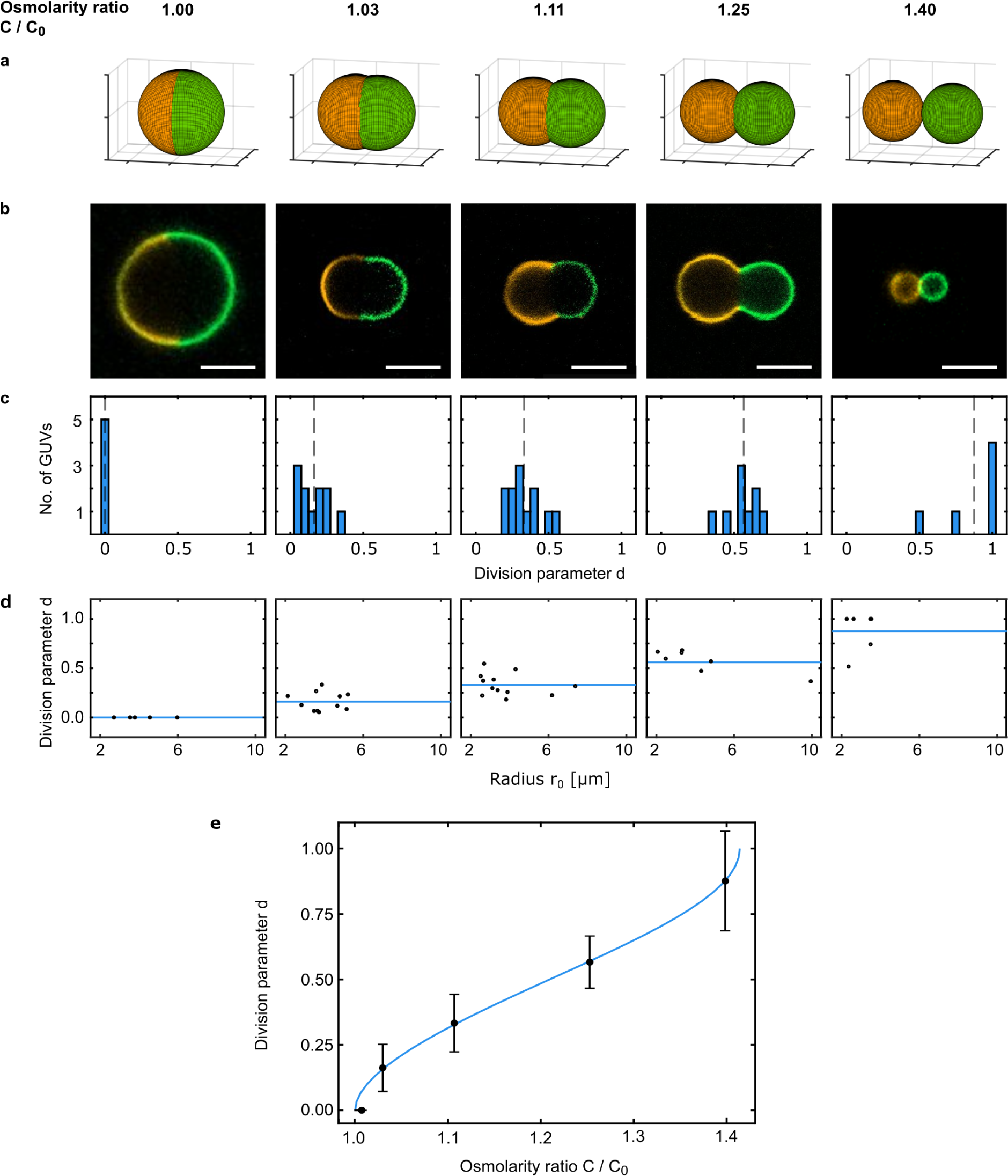
Comparison of experimental results with theoretical predictions of the GUV shape at different osmolarity ratios. **a** Theoretically predicted shapes of symmetric GUVs at different osmolarity ratios *C/C*_0_ as indicated. **b** Representative confocal fluorescence images of symmetric phase-separated GUVs at the corresponding osmolarity ratios *C/C*_0_. The ld phase is labeled with LissRhod PE (orange), the lo phase with 6-FAM-labeled cholesterol-tagged DNA (green). Scale bars: 10 µm. **c** Histograms of division parameter *d* extracted from multiple confocal fluorescence images. Dashed gray lines indicate the mean. **d** Scatter plots of the experimentally determined division parameters plotted against the radius of the initially spherical GUVs. Solid blue lines represent the theoretical prediction, which postulates size-independence of the division process. **e** Division parameter *d* as a function of osmolarity ratio *C/C*_0_. The mean values of the measured division parameters (black) and the theoretical prediction from Equation 1 (solid blue line) are plotted. Error bars correspond to the standard deviation of the values for *d* extracted from independent confocal images.

### Symmetric and asymmetric division regulated by metabolic activity

Any process that leads to an increase of the osmolarity in the outer aqueous phase of phase-separated GUVs should, in principle, trigger their division. Metabolic processes, i.e. the decomposition of molecules through enzymes, inevitably lead to such an osmolarity increase and should thus be suitable to regulate division. We thus set out to metabolize the sugar solution surrounding the GUVs using enzymatic digestion as a trigger for division. Sucrose is decomposed into fructose and glucose by the metabolic enzyme invertase in aqueous solution. The enzymatic reaction is displayed in Figure 4**a**, Figure 4**b** shows the protein structure of invertase with its four subunits [38]. In order to quantitatively analyze the division process, we measure the osmolarity of the GUV-containing solution after addition of invertase over time. Figure 4**c** shows that the osmolarity is indeed increasing over time, confirming the activity of the enzyme. Since the enzyme concentration is directly related to reaction rate, we perform osmolarity measurements at different concentrations of invertase to define suitable conditions for GUV division. We find that 44 mg l^*−*1^ invertase leads to a 1.41-fold increase in the osmotic pressure ratio *C/C*_0_ within 36 min see Figure 4**c** – which should, in turn, lead to the division of symmetric GUVs at this time point. GUVs with an assymmetric lipid composition (*l* ≠ 0.5), however, should divide at earlier time points according to our model.

**Figure 4:**
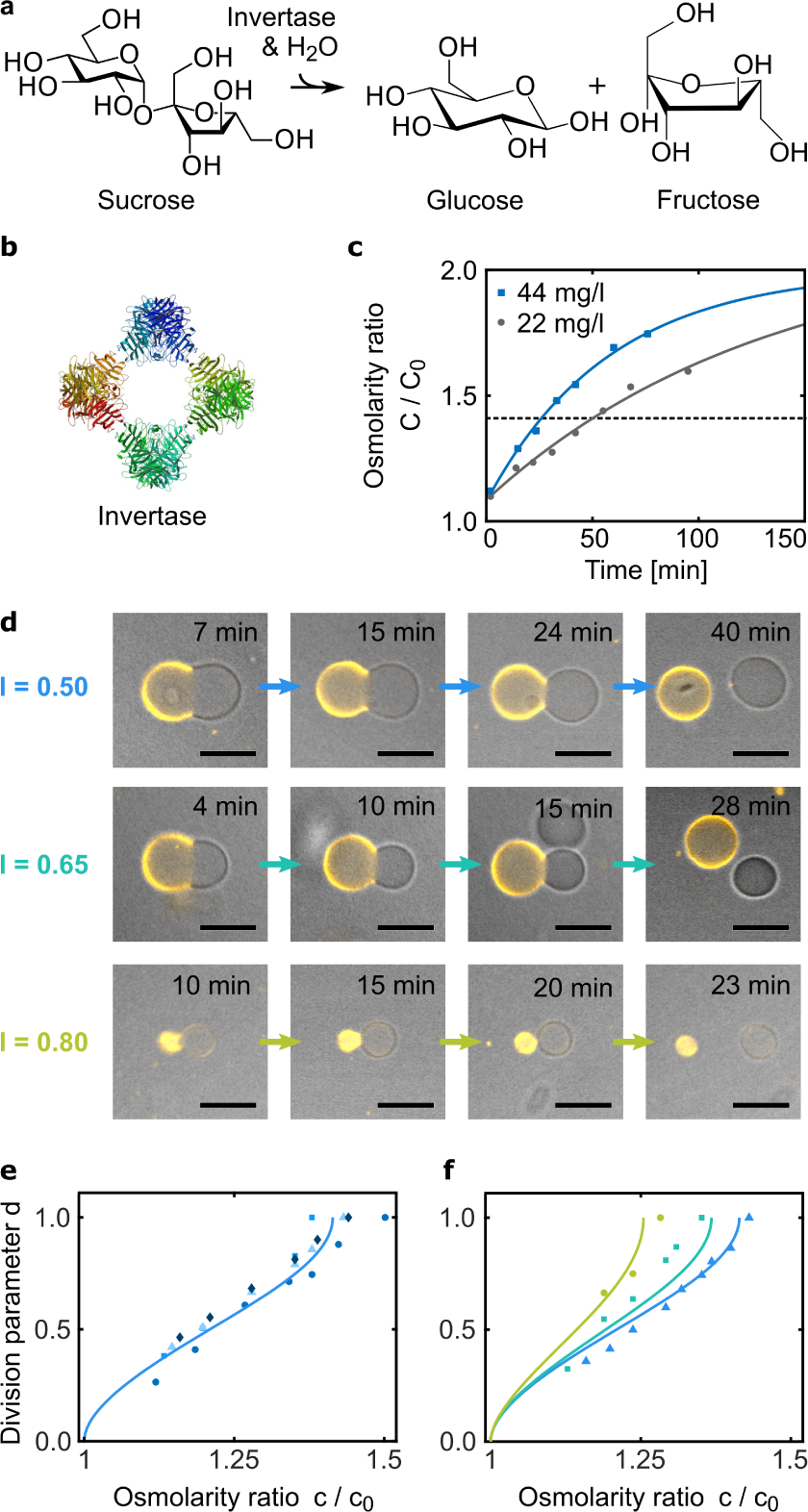
Division of GUVs triggered by metabolic activity. **a** Chemical reaction pathway of sucrose degradation catalyzed by the enzyme invertase. **b** Protein structure of invertase, PDB ID: 4EQV [38]. **c** Osmolarity ratio *C/C*_0_ as a function of time for GUV-containing solutions composed of 300 mM sucrose, 10 mM HEPES and 44 mg l^*−*1^ invertase (Solution 1, blue), or 22 mg l^*−*1^ invertase (gray). Error bars are too small to be visible. The data was fitted with limited growth fits (solid lines). The dotted black line indicates the osmolarity ratio for which division of symmetric GUVs (*l* = 0.5) is expected to occur. **d** Corresponding confocal fluorescence images at different time points during the division process of GUVs with different lipid ratios in the presence of 44 mg l^*−*1^ invertase (Solution 1). The images are overlays of confocal fluorescence (ld phase labeled by LissRhod PE) and brightfield images. Scale bars: 10 µm. **e** Division parameter *d* of four different GUVs with a lipid ratio of *l* = 0.5 in Solution 1 plotted against the osmolarity ratio *C/C*_0_. The values for *C/C*_0_ were obtained from the osmolarity measurements displayed in **c**. The solid blue line shows the theoretically predicted division curve. **f** Division parameter *d* of GUVs with a lipid ratio of *l* = 0.5 (blue), *l* = 0.65 (cyan) and *l* = 0.8 (orange) in Solution 1 plotted against the osmolarity ratio *C/C*_0_. Solid lines are the theoretically predicted division curves for the corresponding lipid ratios.

To test this prediction experimentally, we carry out confocal fluorescence imaging experiments in parallel to the osmolarity measurement. As visible in Figure 4**d**, we can indeed trigger GUV division based on metabolic activity. This confirms once again that the division of phase-separated GUVs relies solely on a change in the osmolarity ratio, as postulated in our model. Notably, the division of the symmetric GUVs (top row) happens after approximately 38 min as predicted. Under the same conditions, assymmetric GUVs exhibit shorter division times, corresponding to lower osmolarity ratios (approximately 27 min for *l* = 0.65 and 20 min for *l* = 0.80, Figure 4**d**, middle and bottom row, respectively). Note that from the osmolarity measurements, we found that Mg^2+^ leads to a reduction of invertase activity in the presence of GUVs (see Supplementary Figure S6). This could be due to interactions between the enzyme and the membrane of the GUVs mediated by divalent ions. Thus, Mg^2+^ was omitted from the buffer. Since Mg^2+^ is necessary for the incorporation of cholesterol-tagged DNA which labels the lo phase (see Supplementary Figure S1), overlays of bright field images and confocal fluorescence images of the ld phase were used. The advantage of performing the division experiment in the absence of Mg^2+^ is that the GUVs are not prone to stick to one another due to electrostatic interactions after the division is completed. Importantly, we can thereby confirm that the division is complete, without a remaining tight neck, for all observed GUVs. To quantify our experimental results in comparison to the theoretical predictions, Figure 4**e** plots the division parameter *d* of symmetric GUVs (*l* = 0.5) over the osmotic pressure ratio *C/C*_0_. All observed GUVs follow the theoretically predicted division curve (solid line). Furthermore, we were able to quantitatively compare the predictions concerning the behavior of GUVs with different lipid ratios *l*. As shown in Figure 4**f**, we find that more asymmetric GUVs undergo a faster division process because they require lower osmotic pressure ratios – again in good agreement with the theoretical predictions (solid lines, Equation 2). We have thus shown that two fundamentally distinct yet highly abundant processes – metabolic activity and water evaporation – can trigger the division of phase-separated GUVs relying on osmosis. By regulating the transcription of a metabolic enzyme like invertase, cells could, in principle, maintain a high level of control over their division without a sophisticated division machinery.

## Discussion

Synthetic cell division is one of the most exciting albeit challenging tasks towards the bottom-up construction of cellular systems. Our study realizes the protein-free division of GUVs, fully controllable by two physical parameters – phase separation and osmosis. Phase separation of the lipids in the GUV membrane defines a division plane such that an increase of the surface-to-volume ratio by osmosis leads to contraction at the phase boundary and thus the formation of two daughter compartments. By predicting the division process and its outcome based on geometrical considerations, we control the duration of the division process as well as the time point of division and the relative size of the daughter compartments. Most remarkably, we show that GUV division can be regulated autonomously by metabolic activity. Growth of GUVs, on the other hand, has already been demonstrated previously [39, 40, 41] – making the implementation of multiple growth and division cycles an exciting next goal. This could, for instance, be realized by targeted fusion [42] of SUVs composed of the opposite lipid type. The integration of information storage and replication will be yet another important milestone towards the transition from matter to life, or, in other words, towards a synthetic cell which truly deserves its name. In the meantime, our engineering approach to synthetic cell division prompts questions about cellular life as we know it: We may be curious to discover whether phase separation and osmosis may have sustained compartment division at the onset of life, possibly regulated by the expression of metabolic enzymes. And we may ask how remnants thereof play a role in cell biology today – continuously nurturing the emergence of cells from preexisting cells.

## Methods

### GUV formation

Atto488-labelled 1,2-Dioleoyl-sn-glycero-3-phosphoethanolamin (Atto488-DOPE) was purchased from ATTO-TEC GmbH. All other lipids were purchased from Avanti Polar Lipids, Inc., and stored in chloroform at *−*20 °C. Giant unilamellar vesicles (GUVs) were produced via the electroformation method [43] using a VesiclePrepPro device (Nanion Technologies GmbH). Two types of GUVs were produced. Phase-separated GUVs composed of 27.125 % 1,2-dioleoyl-sn-glycero-3-phosphocholine (DOPC), 24,75 % cholesterol, 37.125 % 1,2-dipalmitoyl-sn-glycero-3-phosphocholine (DPPC), 10 % cardiolipin (CL), 1 % 1,2-dipalmitoyl-sn-glycero-3-phosphoethanolamine-N-(lissamine rhodamine B sulfonyl) (LissRhod PE) and single-phase GUVs composed of 70 % L-*α*-phosphatidylcholine (EggPC), 29 % L-*α*-phosphatidylglycerol (EggPG), 1 % Atto488-DOPE. 40 µl of 1 mM lipid mix in CHCL_3_ were homogeneously spread on the conductive side of an indium tin oxide (ITO) coated glass coverslide (Visiontek Systems Ltd) using a cover slide. The lipid-coated ITO slide was subsequently placed under vacuum for at least 30 min to achieve complete evaporation of the CHCL_3_. A rubber ring with a diameter of 18 mm was placed on lipid-coated ITO slide. The ring was filled with 275 µl buffer solution, before creating a sealed chamber by placing a second ITO slide on top. The buffer solution used for the phase-separated GUVs was preheated to 65 °C and contained 300 mM sucrose (Sigma-Aldrich Corp.) and 10mM HEPES (Sigma-Aldrich Corp.). For the single-phase GUVs a 500 mM sucrose solution was used as a buffer solution. The assembled electroformation chamber was placed into the VesiclePrepPro and connected to the electrodes. A programmable AC field was applied across the ITO slides. For the phase-separated GUVs, a custom-written multi-step program with defined temperature, voltage, AC-frequency and duration was used (see Table S1). For the single-phase GUVs the preinstalled *Standard* program was selected (see Table S2). GUVs were collected immediately after and stored at 4 °C for up to 2 days.

### Confocal fluorescence microscopy

A confocal laser scanning microscope LSM 800 (Carl Zeiss AG) was used for fluorescence imaging. The images were acquired using a 20x air (Objective Plan-Apochromat 20x/0.8 M27, Carl Zeiss AG) and a 40x water immersion objective (LD C-Apochromat 40x/1.1 W, Carl Zeiss AG). To visualize the phase separation of the GUVs, 6-FAM-labelled cholesterol-tagged DNA (Integrated DNA Technologies, Inc.; DNA sequence: 5’ 6-Fam-CTATGTATTTTGCACAGTTT-Chol 3’; HPLC purified) was used which partitioned mainly into the lo phase [33]. LissRhod PE labelled the ld phase. 6-FAM and Atto488-DOPE were excited with a 488 nm laser diode (Carl Zeiss AG), LissRhod PE with a laser diode at 561 nm (Carl Zeiss AG). Images were analyzed with ImageJ (contrast and brightness adjustments and TrackMate) and Matlab.

### Theoretical predictions

All calculations were carried out with MathWorks Matlab (9.5.0.944444 R2018b) and Jupyter (v. 4.4.0) as described in the main text.

### Determination of osmolality

The osmolality of all solutions was measured with the Osmomat 030 (Gonotec GmbH). Before use, the osmometer was calibrated with calibration solutions of 0, 300 and 900 mOsm/kg (Gonotec GmbH). Each measurement was carried out with a sample volume of 50 µl. Note that for the quantities that are calculated here, the osmolality is a good approximation for the osmolarity (see Supplementary Note). The measurement error of the osmometer itself (in terms of reproducibility) is below 0.5%.

### Enzymatic osmolarity change

Invertase from bakers’s yeast (S. cerevisiae) grade VII, *≥* 300 units/mg was purchased from Sigma-Aldrich Corp. Nominally, one unit of the enzyme hydyrolyzes 1 µmol of sucrose per minute to produce fructose and glucose at pH 4.5 at 55 °C. The GUV solution obtained from electroformation was mixed with a solution containing 1 mg ml^*−*1^ invertase such that the final concentration of invertase was 44.4 mg l^*−*1^ and in a second experiment 22.2 mg l^*−*1^. Immediately after mixing, a part of the solution was used for observation by confocal fluorescence microscopy in a sealed observation chamber. The other part was used for osmolality measurements over time. The experiments were carried out at room temperature.

## Acknowledgments

The authors acknowledge funding from the European Research Council, Grant Agreement no. 294852, SynAd and the MaxSynBio Consortium, which is jointly funded by the Federal Ministry of Education and Research of Germany and the Max Planck Society. They also acknowledge the support from the SFB 1129 of the German Science Foundation and the VolkswagenStiftung (priority call ‘Life?’). J.P.S. is the Weston Visiting Professor at the Weizmann Institute of Science and part of the excellence cluster CellNetworks at the University of Heidelberg. K.G. received funding from the European Union’s Horizon 2020 research and innovation program under the Marie Skłodowska-Curie grant agreement No 792270. The Max Planck Society is appreciated for its general support.

## Author contributions

All authors contributed extensively to the work presented here. Y.D. performed the experiments, J.P.S. and K.G. supervised the research, Y.D. and K.G wrote the manuscript.

## Additional information

**Supplementary information** is available for this paper.

**Correspondence and requests for materials** should be addressed to K.G.

## Data availability

All data is available in the main text or the Supplementary information.

