## Supplementary information for "Protein-free division of giant unilamellar vesicles controlled by enzymatic activity"

**3 Supplementary Note: Osmolarity vs. osmolality** **12**

**Supplementary References** **14**

### 1 Supplementary Tables

#### 1.1 Table S1: Electroformation protocol for phase-separated GUVs

| **Step** | **Time [s]** | **Ampl [V]** | **Freq [Hz]** | **Temp [°C]** |
|---|---|---|---|---|
| Initiate | 300 | 1 | 10 | 70 |
| Main | 2100 | 1 | 10 | 70 |
| Detach1 | 2160 | 1 | 3 | 70 |
| Detach2 | 3000 | 1 | 3 | 70 |
| Detach3 | 3060 | 1 | 1 | 70 |
| Detach4 | 3420 | 1 | 1 | 70 |
| Detach5 | 3480 | 1 | 0.5 | 70 |
| Detach6 | 3840 | 1 | 0.5 | 70 |
| Detach7 | 3960 | 1 | 0 | 70 |

**Supplementary Table S1**
Electroformation protocol for the formation of phase-separated GUVs using the Vesicle Prep Pro (Nanion Technologies GmbH). Custom-written multi-step program for formation of phase-separated GUVs, adapted from a previously published protocol [1]. Note that it is crucial to keep the temperature above the phase-transition temperature throughout the protocol. Parameters are changed linearly over time from one step to the next.

#### 1.2 Table S2: Electroformation protocol for conventional GUVs

| **Step** | **Time [s]** | **Ampl [V]** | **Freq [Hz]** | **Temp [°C]** |
|---|---|---|---|---|
| Initiate | 180 | 3 | 5 | 37 |
| Main | 7380 | 3 | 5 | 37 |
| Detach | 7680 | 0 | 5 | 37 |

**Supplementary Table S2**
Electroformation protocol for single-phase GUV formation using the Vesicle Prep Pro (Nanion Technologies GmbH). The programme was preinstalled as the standard protocol for GUV formation. Parameters are changed linearly over time from one step to the next.

### 2 Supplementary Figures

#### 2.1 Figure S1: Necessity of $MgCl_2$ for attachment of cholesterol-tagged DNA

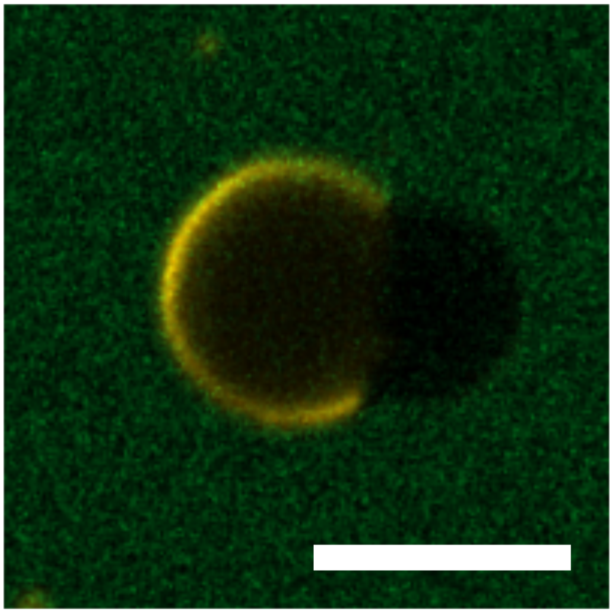

**Supplementary Figure S1**
Confocal fluorescence microscope image of a GUV (ld phase labeled with LissRhod PE, orange) in a solution of 300 mM sucrose, 10 mM HEPES and 1 µM 6-FAM-labeled cholesterol-tagged DNA (green). Due to lack of $MgCl_2$, the cholesterol-tagged DNA does not attach to the GUV membrane and is homogeneously distributed in the outer aqueous phase instead. For this reason, 10 mM $MgCl_2$ was added to the outer aqueous phase for Figures 1 and 3 (main text), resulting in the attachment of the cholesterol-tagged DNA to the lo phase. Since $MgCl_2$ inhibited the activity of invertase (see Figure S5), it was not possible to label the lo phase in Figure 4 (main text). Scale bar: 10 µm

.

#### 2.2 Figure S2: $MgCl_2$-mediated electrostatic interaction between divided GUVs may be overcome through gentle shaking.

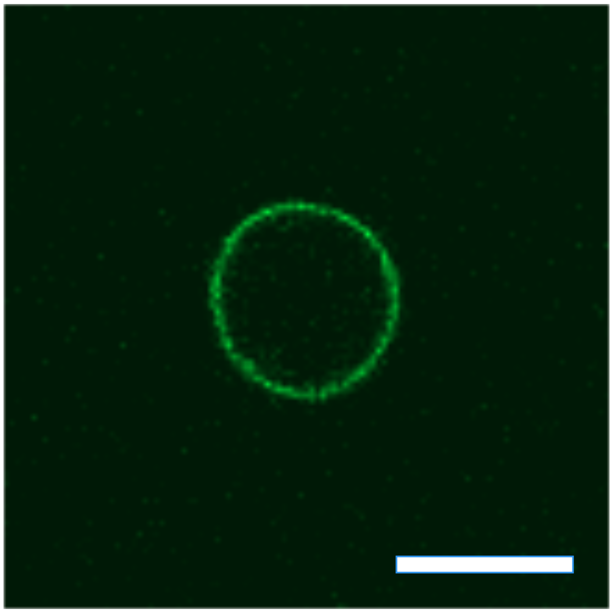

**Supplementary Figure S2**
Confocal fluorescence microscope image of a GUV consisting only of the lo phase (green, 6-FAM labeled cholesterol-tagged DNA partitioned into the lo phase) after mixing with a higher concentrated sucrose solution leading to an osmolarity ratio of $C/C_0 = 1.44$. At this ratio all GUVs are fully divided, yet often adhere to one another if $Mg^{2+}$ is present in the buffer (see Figure 1, main text). Observation of the mixture after gentle shaking, however, yields a high amount of single-phased GUVs. A possible explanation is that $Mg^{2+}$-mediated electrostatic interactions between divided GUVs could be overcome mechanically due to the mixing process. Scale bar: 10 µm.

#### 2.3 Figure S3: Lipid tubulation caused by osmotic pressure

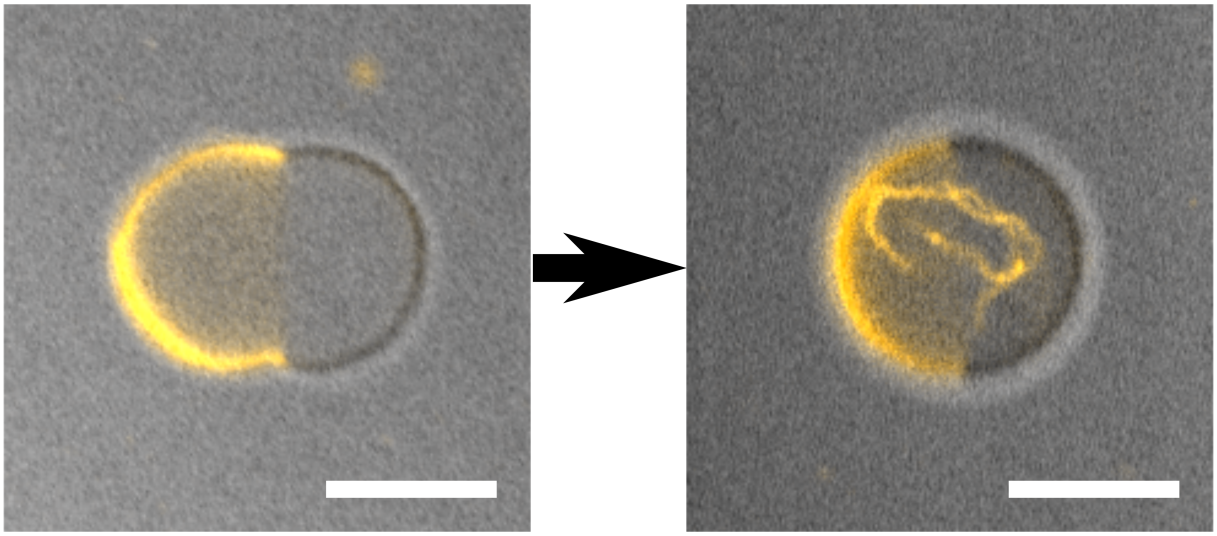

**Supplementary Figure S3**
Lipid tubulation as a result of fast changes in osmotic pressure. Left and right: Overlay of confocal fluorescence (ld phase labeled with LissRhod PE) and bright field images of a GUV exposed to an osmolarity gradient ($C > C_0$). Water efflux leads to a reduction of the GUV's inner volume. The GUV deforms initially (left image). As the osmotic pressure increases over time, lipid tubulation is observed and the spherical shape is restored (right image). This effect inhibits successful division of the GUVs. It occurred mainly when the osmolarity is too high or changes too quickly. Scale bars: 10 µm.

#### 2.4 Figure S4: Osmolarity mismatch after electroformation

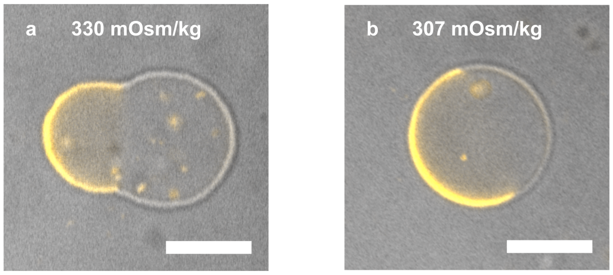

**Supplementary Figure S4**

Overlays of confocal fluorescence (ld phase labeled with LissRhod PE) and bright field images of GUVs in buffers of different osmolality. (**a**) Directly after electroformation, the GUVs exhibit a non-spherical shape even though the measured osmolality of the vesicle-containing solution does not increase enough for such a significant shape change ($325\,\mathrm{mOsm\,kg^{-1}}$ before electroformation, $330\,\mathrm{mOsm\,kg^{-1}}$). (**b**) If the buffer solution is diluted to $307\,\mathrm{mOsm\,kg^{-1}}$, the GUV returns to its spherical shape. This may be due to the fact that the vesicles still grow after forming a sealed compartment, leading to a reduced sucrose concentration inside the vesicle. We thus diluted the outer aqueous phase until the GUVs were spherical to achieve our desired initial conditions. Scale bars: 10 µm.

#### 2.5 Figure S5: Microfluidic trapping approach

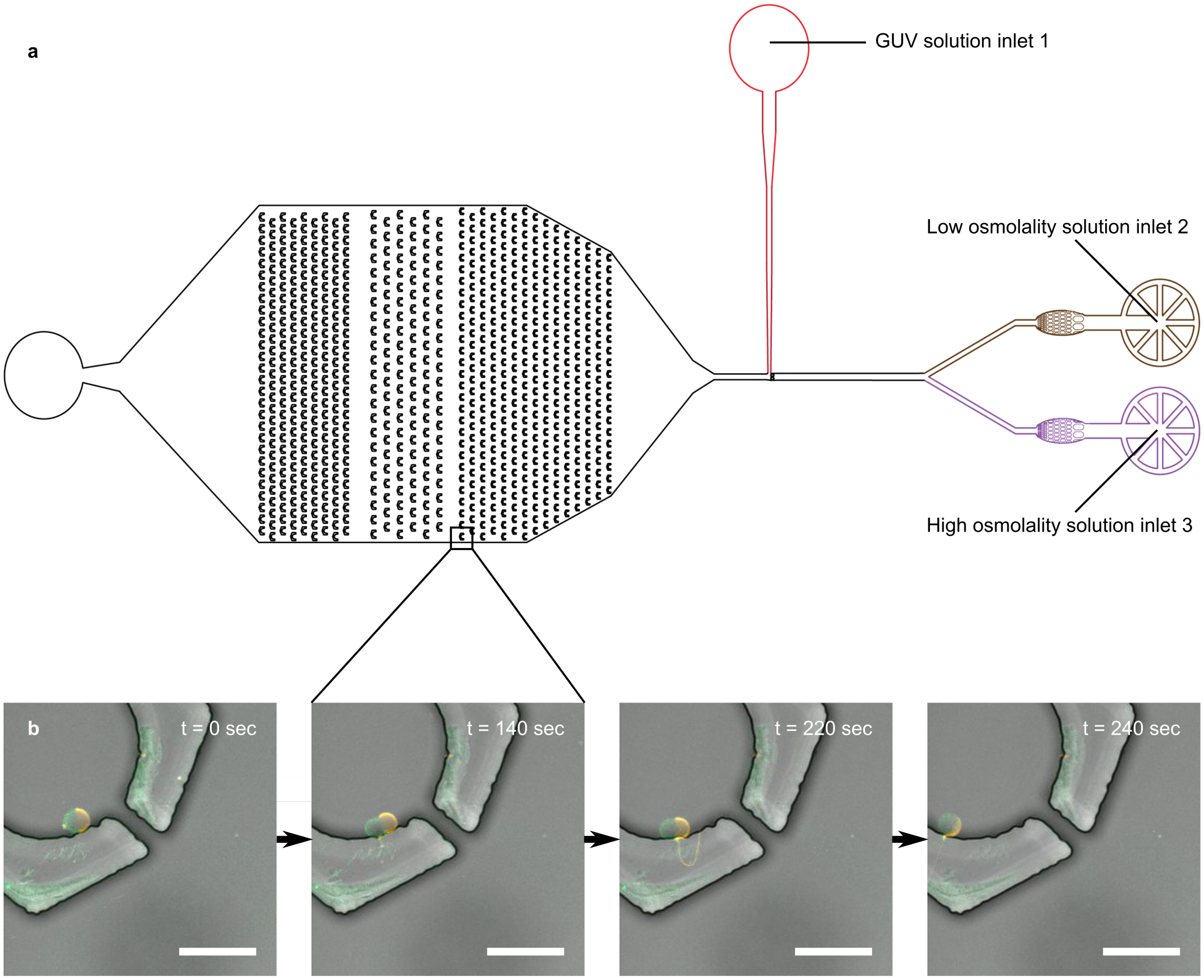

**Supplementary Figure S5**
(Continued on the following page.)

**Supplementary Figure S5**

Microfluidic approach for trapping and observation of GUV division. **a** Sketch of the microfluidic trapping device used for trapping of phase-separated GUVs. First, a solution containing GUVs was flushed into the device via Inlet 1. Subsequently, a low osmolality solution ($280\,\mathrm{mOsm\,kg^{-1}}$, same osmolality as within the GUVs) was flushed into the device at a constant flow rate of $1\,\mu\mathrm{l/\,min}$ via Inlet 2. In order to gradually increase the effective osmolality around the GUVs, a second high osmolality solution ($600\,\mathrm{mOsm\,kg^{-1}}$) was flushed in via Inlet 2, starting at a flow rate of $0\,\mu\mathrm{l/\,min}$ which was gradually increased to $2\,\mu\mathrm{l/\,min}$ over 40 min. **b** Time series of overlays of confocal fluorescence (ld phase labeled with LissRhod PE and lo phase labeled with 6-FAM-labeled cholesterol-tagged DNA) and bright field images of a phase-separated GUV in a trapping device. The interaction of the GUVs with the coverslide and the PDMS microstructures can lead to effects that inhibit the division process. The deformation process can be altered through contact of the GUV with solid interfaces and lead to lipid tubulation rather than splitting as visible in the confocal images. Scale bars: $30\,\mu\mathrm{m}$.

#### 2.6 Figure S6: Reduction of invertase activity in the presence of $MgCl_2$

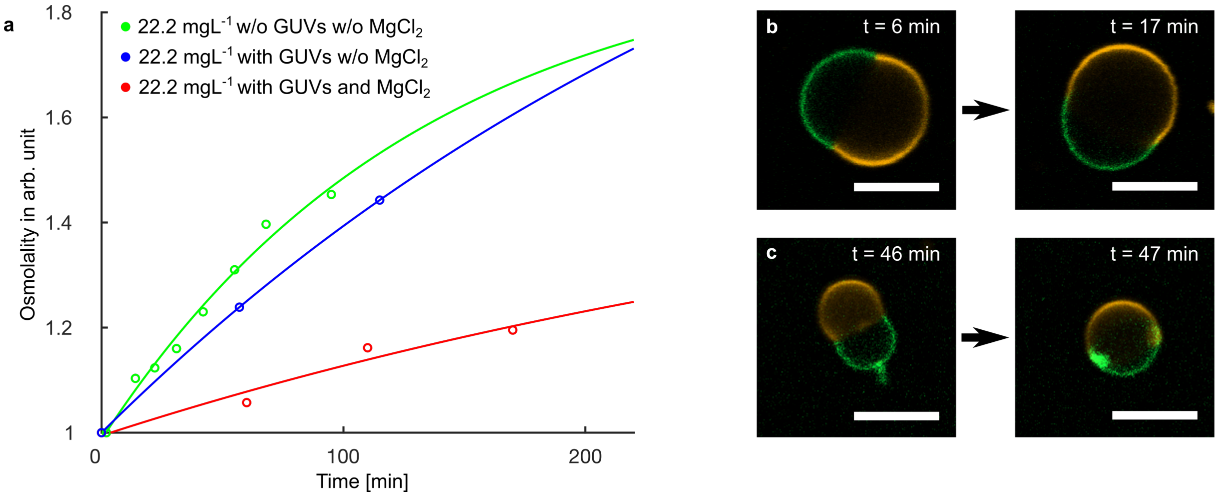

**Supplementary Figure S6**
Effect of $MgCl_2$ on invertase activity in the presence of GUVs. **a** Normalized osmolality measurements over time of a buffer containing 300 mM sucrose, 10 mM HEPES and $22.2\,\mathrm{mg\,l^{-1}}$ in the absence of GUVs and $MgCl_2$ (green); with GUVs but without $MgCl_2$ (blue) and with 10 mM $MgCl_2$ and GUVs (red). Solid lines are limited growth fits. We hypothesize that the reduction of invertase activity in the presence of $MgCl_2$ and GUVs could be caused by a charge-mediated adhesion of the invertase to the GUVs. **b** + **c** Confocal fluorescence microscopy images of GUVs in the presence of invertase and $MgCl_2$. The ld phase is labeled by LissRhod PE (orange) and the lo phase is visualized by 6-FAM-labeled cholesterol-tagged DNA (green). The time after mixing with invertase is indicated. Under these conditions, almost all GUVs did not divide and returned to a spherical shape instead. These observations again point towards an interaction between the invertase and the surface of the GUVs, leading to the reduction of invertase activity. We hence performed the invertase experiments in the absence of $MgCl_2$.

### 3 Supplementary Note: Osmolarity vs. osmolality

It is important to note that the measurements produced by the Osmomat 030 (Genotec GmbH) indicate the osmolality $b$ of the sample solution. However, according to van't Hoffs law the osmotic pressure depends on the total particle concentration in the solution (osmolarity) $C$: $\Pi = CRT$, where $C$ is the osmolarity of the solution, $R$ the ideal gas constant and $T$ the temperature. The osmolarity depends linearly on the osmolality $C = (\rho_S - c_a) \cdot b$ where $\rho_S$ is the density of the solution and $c_a$ is the anhydrous solute concentration. The prefactor $(\rho_S - c_a)$ is dependent on the solute. It can be neglected in most cases since $(\rho_S - c_a) \approx 1\,\mathrm{kg\,l^{-1}}$ [2]. However, for high concentrations of sugars as have been used here, the prefactor can deviate from $1\,\mathrm{kg\,l^{-1}}$. Since we look at the osmotic pressure ratio $C/C_0$, the effect cancels out nevertheless. It can only play a role when using different solutions for the inner and outer phase like in Figure 3 (main text). However, even then, the deviation of the osmotic pressure ratio is negligible: For the solutions that were used here, $(\rho_S - c_a)$ did not deviate more than 15% from $1\,\mathrm{kg\,l^{-1}}$. Hence, the deviation of the osmotic pressure ratio $C/C_0$ is below 1%. Furthermore, Moser and Frazer suggested the that the osmotic pressure is better described by $\Pi = \frac{n}{V'}RT = \rho_{\mathrm{w}} bRT$, for higher concentrations of glucose or sugar with $V'$ the volume of pure water in the solution, $\rho_{\mathrm{w}}$ is the density of pure water and $b$ is the osmolality of the solution [3]. Therefore, we used the osmolality instead of the osmolarity for the calculations.

Note that within the confocal time series, other free-floating GUVs appear in the field of view, which show similar deformation stages as the GUV that was tracked.
